# AYbRAH: a curated ortholog database for yeasts and fungi spanning 600 million years of evolution

**DOI:** 10.1101/237974

**Authors:** Kevin Correia, Shi M. Yu, Radhakrishnan Mahadevan

## Abstract

Budding yeasts inhabit a range of environments by exploiting various metabolic traits. The genetic bases for these traits are mostly unknown, preventing their addition or removal in a chassis organism for metabolic engineering. To help understand the molecular evolution of these traits in yeasts, we created Analyzing Yeasts by Reconstructing Ancestry of Homologs (AYbRAH), an open-source database of predicted and manually curated ortholog groups for 33 diverse fungi and yeasts in Dikarya, spanning 600 million years of evolution. OrthoMCL and OrthoDB were used to cluster protein sequence into ortholog and homolog groups, respectively; MAFFT and PhyML were used to reconstruct the phylogeny of all homolog groups. Ortholog assignments for enzymes and small metabolite transporters were compared to their phylogenetic reconstruction, and curated to resolve any discrepancies. Information on homolog and ortholog groups can be viewed in the AYbRAH web portal (https://kcorreia.github.io/aybrah/) to review ortholog groups, predictions for mitochondrial localization and transmembrane domains, literature references, and phylogenetic reconstructions. Ortholog groups in AYbRAH were compared to HOGENOM, KEGG Orthology, OMA, eggNOG, and PANTHER. PANTHER and OMA had the most congruent ortholog groups with AYbRAH, while the other phylogenomic databases had greater amounts of under-clustering, over-clustering, or no ortholog annotations for proteins. Future plans are discussed for AYbRAH, and recommendations are made for other research communities seeking to create curated ortholog databases.

## 1 INTRODUCTION

Yeasts are unicellular fungi that exploit diverse habitats on every continent, including the gut of wood boring beetles, insect frass, tree exudate, rotting wood, rotting cactus tissue, soil, brine solutions, and fermenting juice (Kurtzman et al., 2011). The most widely studied yeasts are true budding yeasts, which span roughly 400 million years of evolution in the subphylum Saccharomycotina (Hedges et al., 2015), and possess a broad range of traits important to metabolic engineering. These include citrate and lipid accumulation in *Yarrowia* (Aiba and Matsuoka, 1979) and *Lipomyces* (Boulton and Ratledge, 1983), thermotolerance in multiple lineages (Banat et al., 1996; Ryabova et al., 2003), acid tolerance in *Pichia* (Rush and Fosmer, 2013) and *Zygosaccharomyces* (Lindberg et al., 2013), methanol utilization in *Komagataella* (Hazeu and Donker, 1983), osmotolerance in *Debaryomyces* (Larsson and Gustafsson, 1993), xylose to ethanol fermentation in multiple yeast lineages (Schneider et al., 1981; Slininger et al., 1982; Toivola et al., 1984), alternative nuclear codon assignments (Mühlhausen et al., 2016), glucose and acetic acid co-consumption in *Zygosaccharomyces* (Sousa et al., 1998), and aerobic ethanol production (the Crabtree effect) in multiple lineages (de Deken, 1966; van Urk et al., 1990; Blank et al., 2005; Christen and Sauer, 2011). The complete genetic bases of these traits are mostly unknown, preventing their addition or removal in a chassis organism for biotechnology.

The distinction between orthologs, paralogs, ohnologs and xenologs plays an important role in bridging the genotype-phenotype gap across the tree of life (Koonin, 2001). Briefly, orthologs are genes that arise from speciation and *typically* have a conserved function; paralogs and ohnologs emerge from locus and whole genome duplications, respectively, and *may* have a novel function; xenologs derive from horizontal gene transfer between organisms (Jensen, 2001; Koonin, 2005). Knowledge of these types of genes has played an important role in deciphering *Saccharomyces cerevisiae*’s physiology. For example, the Adh2p paralog in *S. cerevisiae* consumes ethanol and evolved from an ancient Adh1p duplication whose kinetics favoured ethanol production (Thomson et al., 2005); the whole genome duplication led to the *MPC2* and *MPC3* ohnologs in the *Saccharomyces* genus, which encode the fermentative and respirative subunits of the mitochondrial pyruvate carrier (Bender et al., 2015), respectively; the *URA1* xenolog from Lactobacillales enables uracil to be synthesized anaerobically in most Saccharomycetaceae yeasts (Hall et al., 2005). These examples demonstrate how understanding the origin of genes has narrowed the genotype-phenotype gap for fermentation in Saccharomycetaceae.

Many genomics studies have focused on the Saccharomycetaceae family, and to a lesser extent the CTG clade (Dujon, 2010), but more can be learned about yeast metabolism by studying its evolution over a longer time horizon, especially with yeasts having deeper phylogeny (Gaucher et al., 2010). If we could study the metabolism of the mother of all budding yeasts, which we refer to as the Proto-Yeast, we could track the gains and losses of orthologs and function in all of her descendants to bridge the genotype-phenotype gap. Proto-Yeast has evolved from her original state, making this direct study impossible, but we can reconstruct her metabolism through her living descendants. In recent yeasts, dozens of yeasts with deep phylogeny have been sequenced (Riley et al., 2016), paving the way for greater insight into the evolution of metabolism in yeasts beyond Saccharomycetaceae.

Ortholog databases are critical to facilitating comparative genomics studies and inferring protein function. Most of these databases are constructed using graph-based methods that rely on sequence similarity, while fewer databases use tree-based methods (Kuzniar et al., 2008). Existing ortholog databases do not span diverse yeasts (Figure 1), and sometimes cannot distinguish between orthologs and paralogs (Tables S1 and S2). In addition to these databases, orthologs are identified on an *ad hoc* basis with OrthoMCL for comparative genomics studies (Wohlbach et al., 2011; Papini et al., 2012), or with the reciprocal best hit method for genome-scale network reconstructions (Caspeta et al., 2012); these orthology relationships often lack transparency or traceability, and therefore cannot be scrutinized or continuously improved by research communities. To solve these outlined problems, and ultimately improve our understanding of budding yeast physiology, we present Analyzing Yeasts by Reconstructing Ancestry of Homologs (AYbRAH) (Figure 2). AYbRAH, derived from the Hebrew name Abra, mother of many, is an open-source database of predicted and manually curated orthologs, their function, and origin. The initial AYbRAH database was constructed using OrthoMCL and OrthoDB. PhyML was used to reconstruct the phylogeny of each homolog group. AYbRAH ortholog assignments for enzymes and small metabolite transporters were compared against their phylogenetic reconstruction and curated to resolve any discrepancies. We discuss the information available in the AYbRAH web portal (https://kcorreia.github.io/aybrah/), issues that arose from reviewing the accuracy of ortholog predictions, compare AYbRAH to established phylogenomic databases, discuss the benefits of open-source ortholog databases, future directions for AYbRAH, and offer recommendations to research communities looking to develop ortholog databases for other taxa.

**Figure 1.**
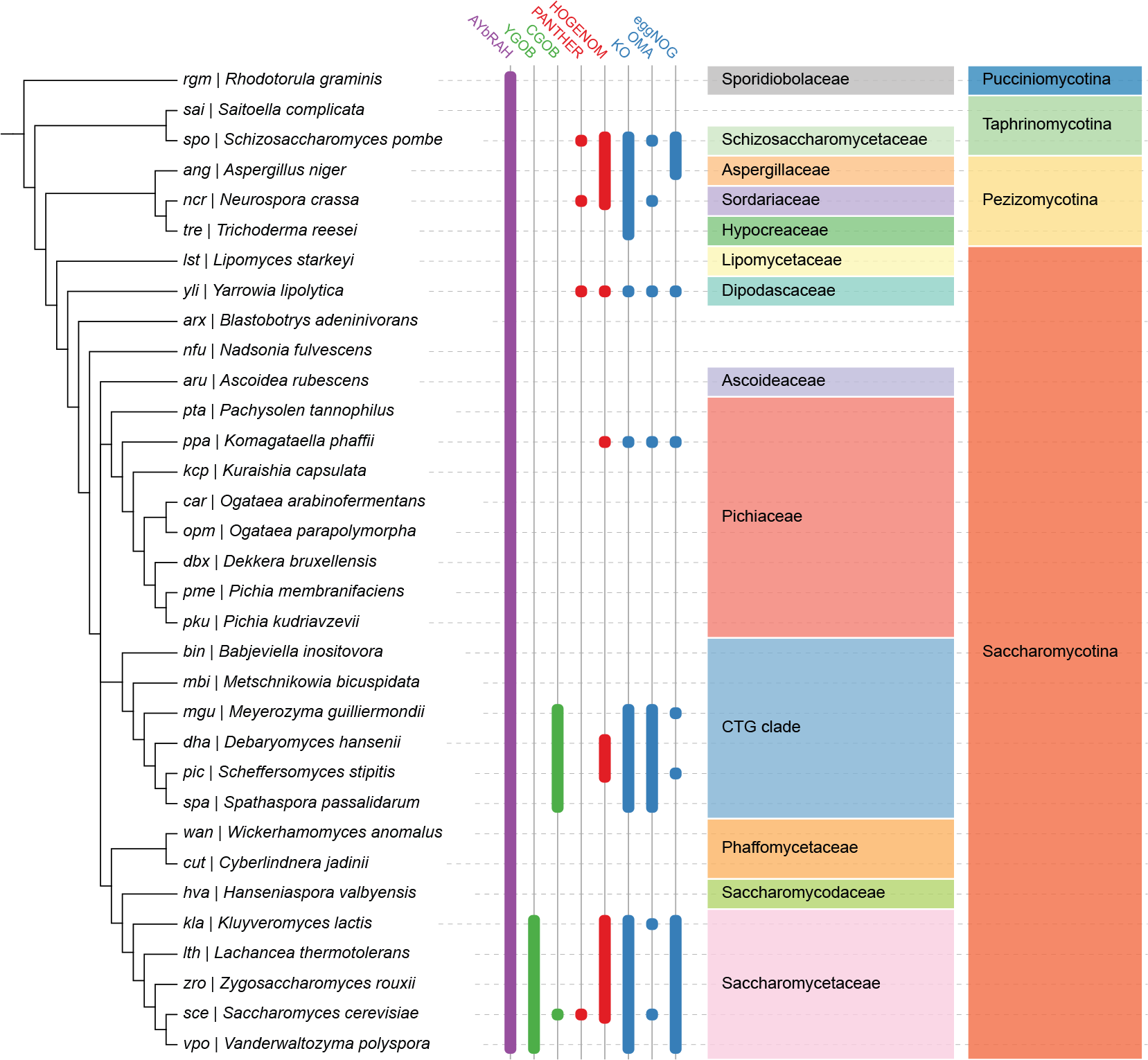
Ortholog database coverage for fungal and yeast genomes in AYbRAH, YGOB, CGOB, PANTHER, HOGENOM, KO, OMA, and eggNOG. Ortholog assignments based on the manual curation of sequence similarity and synteny are shown in green columns; tree-based methods in red columns; graph-based methods in blue columns; a hybrid graph and tree-based method in the purple column. Many ortholog databases are well represented in Saccharomycetaceae and the CTG clade, which had their genomes sequenced during the 2000’s (Dujon, 2010). AYbRAH has ortholog assignments for species in Pichiaceae, Phaffomycetaceae and several *incertae sedis* families, which are not well represented in other ortholog databases, as these yeasts were recently sequenced (Riley et al., 2016). The well established phylogenomic databases span other yeast species not shown in this phylogeny, but they mostly belong to Saccharomycetaceae or the CTG clade.

**Figure 2.**
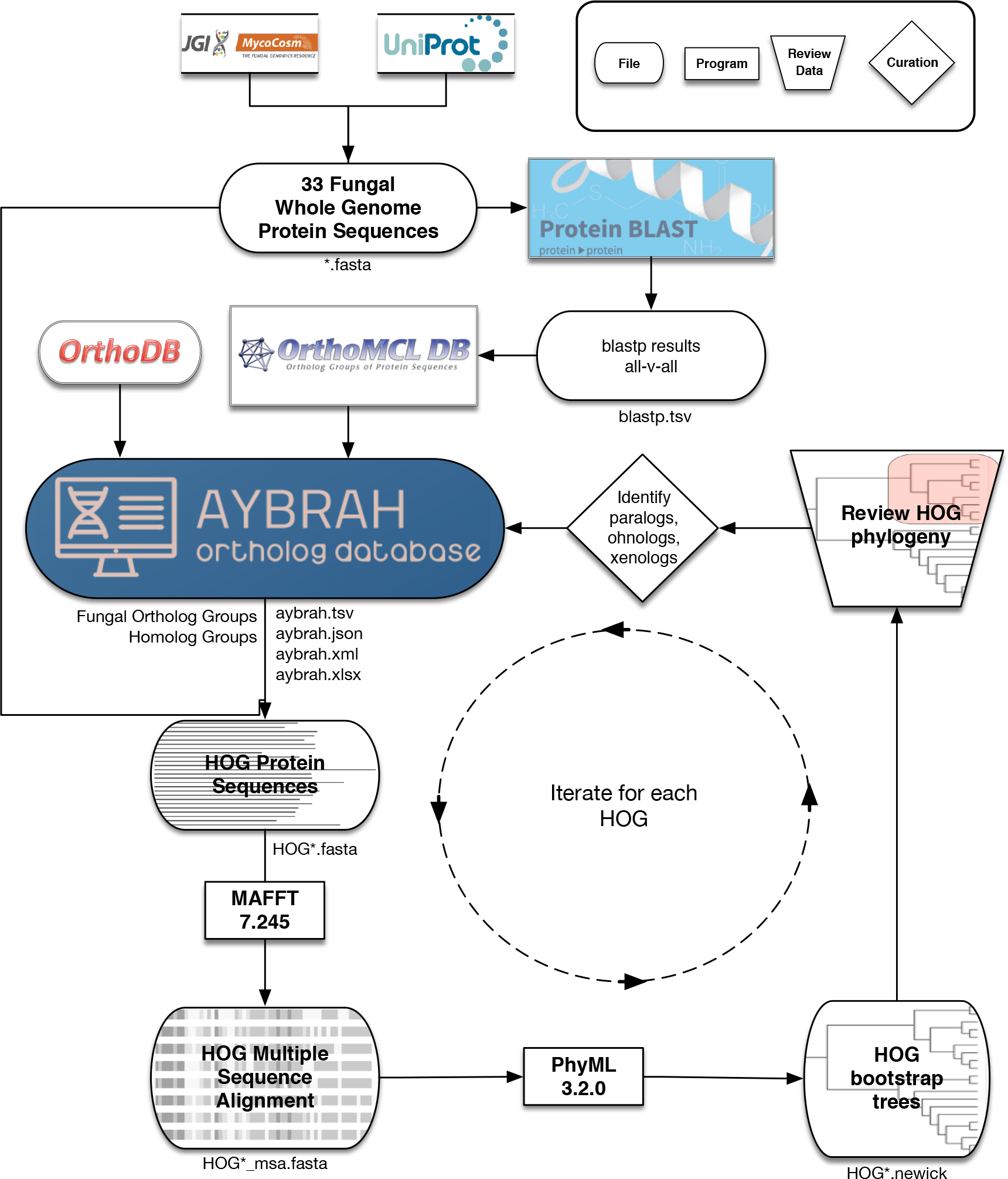
AYbRAH workflow for ortholog curation. 33 fungal and yeast proteomes were downloaded from UniProt and MycoCosm. BLASTP computed the sequence similarity between all proteins. OrthoMCL clustered the proteins into putative Fungal Ortholog Groups (FOG’s) using the BLASTP results. FOG’s were clustered into HOmolog Groups (HOG’s) using Fungi-level homolog assignments from OrthoDB. Multiple sequence alignments for each homolog group were obtained with MAFFT, and 100 bootstrap phylogenetic trees were reconstructed with PhyML. The consensus phylogenetic trees for enzymes and transporters were reviewed and curated to differentiate between orthologs, paralogs, ohnologs, and xenologs.

## 2 METHODS

### Initial construction of AYbRAH

AYbRAH was created by combining several algorithms and databases in a pipeline (Figure 2). 212 836 protein sequences from 33 organisms (Table 1) in Dikarya were downloaded from UniProt (Consortium, 2014) and MycoCosm (Grigoriev et al., 2013). OrthoMCL (Fischer et al., 2011) clustered protein sequences into putative Fungal Ortholog Group’s (FOG’s); default parameters were used for BLASTP and OrthoMCL. The FOG’s from OrthoMCL were coalesced into HOmolog Groups (HOG’s) using Fungi-level homolog group assignments from OrthoDB v8 (Kriventseva et al., 2015).

**Table 1.**
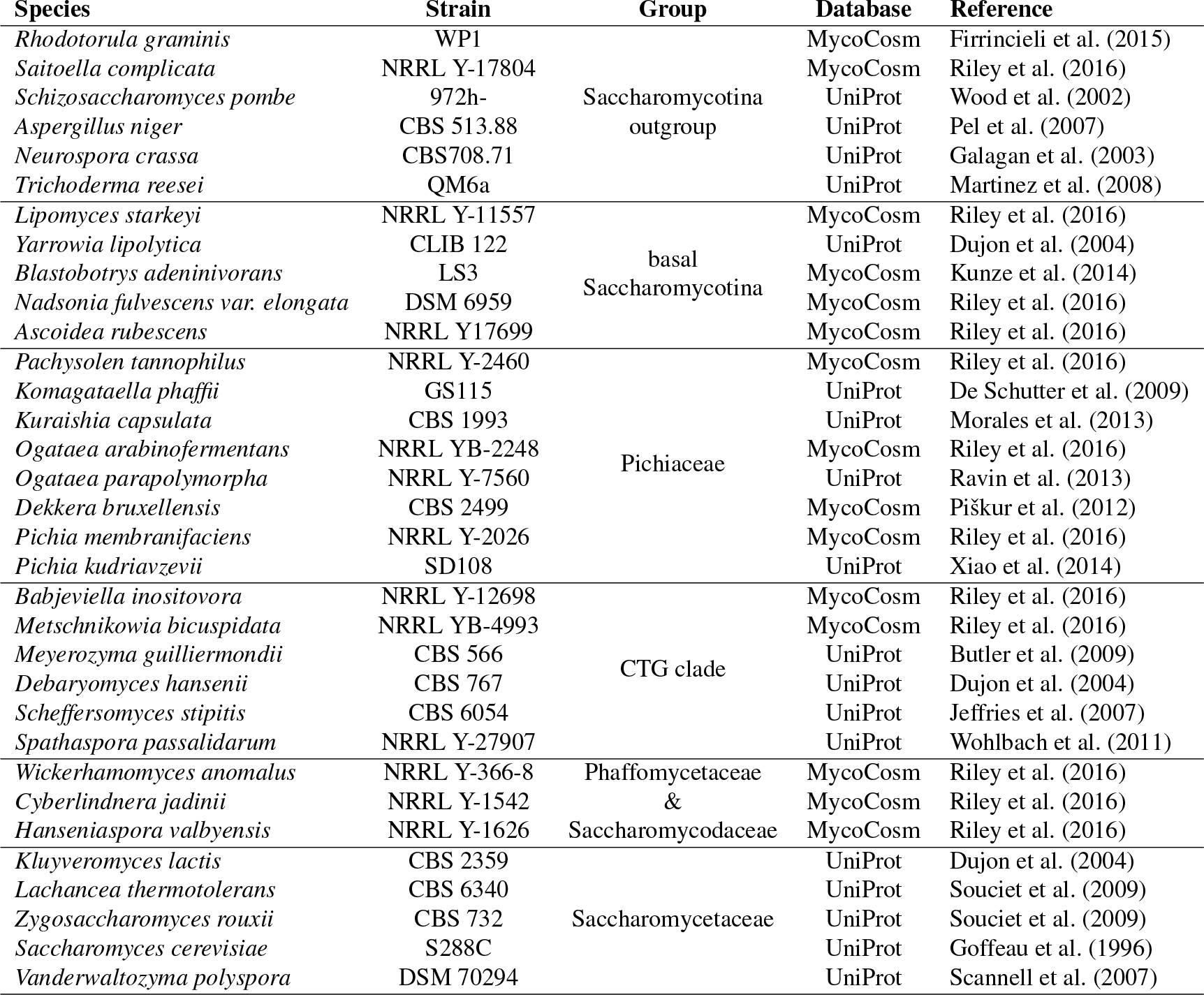
Fungal and yeast strain genomes in AYbRAH. Protein sequences were downloaded from UniProt or MycoCosm. Species were assigned to monophyletic or paraphyletic groups based on divergence time with *Saccharomyces cerevisiae*.

### AYbRAH curation

Multiple sequence alignments were obtained for each homolog group with MAFFT v7.245 (Katoh and Standley, 2013) using a gap and extension penalty of 1.5. 100 bootstrap trees were reconstructed for each HOG with PhyML v3.2.0 (Guindon et al., 2010), optimized for tree topology and branch length. Consensus phylogenetic trees were generated for each HOG with SumTrees from DendroPy v4.1.0 (Sukumaran and Holder, 2010), and trees were rendered with ETE v3 (Huerta-Cepas et al., 2016a). The phylogenetic reconstruction for enzymes and metabolite transporters were reviewed when OrthoMCL failed to differentiate between orthologs and paralogs, caused by over-clustering (Figure 5), or when orthologous proteins were dispersed into multiple ortholog groups, caused by under-clustering. Orthologs were identified by visual inspection of the phylogenetic trees or with an ETE 3-based script (Huerta-Cepas et al., 2016a).

### Annotating additional proteins

Additional steps were required to assign proteins to ortholog groups because OrthoMCL did not cluster all related proteins to ortholog groups, or because whole genome protein annotations were incomplete. First, proteins in OrthoDB homolog groups were added to new FOG’s if they were not assigned to any FOG by OrthoMCL. Next, each organism had its genome nucleotide sequence queried by with a protein of the species closest relative from each FOG using TBLASTN (expect threshold of 1e-20). Annotated proteins were then queried against the TBLASTN hits to determine which proteins were annotated but not assigned to a FOG by OrthoMCL, and which proteins were unannotated despite a match in its nucleotide sequence. Proteins identified via TBLASTN with a sequence length less than 75% of the mean FOG sequence length were discarded from the candidate list. The remaining proteins were assigned to a HOG by its best hit via BLASTP, and to a FOG with pplacer (Matsen et al., 2010) via the MAFFT add alignment option. For example, Cybja1 169606 (A0A1E4RV95), which encodes NADP-dependent isocitrate dehydrogenase in *Cyberlindnera jadinii*, was not assigned to any ortholog group by OrthoMCL despite its high similarity to other proteins. It was added to FOG00618 by pplacer (Matsen et al., 2010) with high confidence. No 60S ribosomal protein L6 (FOG00006) was present in *Meyerozyma guilliermondii*’s protein annotation; it was identified by TBLASTN, annotated as mgu AYbRAH 00173, and added to FOG00006 by pplacer (Matsen et al., 2010).

### Comparison of ortholog groups

AYbRAH ortholog assignments were compared to OMA (Altenhoff et al., 2015), PANTHER (Mi et al., 2016), HOGENOM (Penel et al., 2009), eggNOG (Huerta-Cepas et al., 2016b), KEGG Orthology (Mao et al., 2005). Phylogenomic annotations were downloaded from UniProt. Ortholog groups were assessed as congruent, over-clustered, under-clustered, over and under-clustered, or no ortholog assignment relative to AYbRAH. AYbRAH ortholog groups were only compared with an ortholog database if an ortholog group in AYbRAH had proteins from species present in the other ortholog database. For example, FOG19691 consists of proteins from *Ascoidea rubescens*, *Pachysolen tannophilus*, *Kuraishia capsulata*, *Ogataea parapolymorpha*, *Dekkera bruxellensis*, *Pichia kudriavzevii*, *Pichia membranifaciens*, *Babjeviella inositovora*, *Wickerhamomyces anomalus*, and *Cyberlindnera jadinii*. None of the phylogenomic databases have ortholog assignments for these organisms, and therefore cannot be compared with AYbRAH. Evolview v2 (He et al., 2016) was used to map ortholog databases coverage onto the yeast species tree.

### Subcellular localization prediction

Subcellular localization predictions for all proteins in the pan-genome were computed with MitoProt II (Claros and Vincens, 1996), Predotar (Small et al., 2004), and TargetP (Emanuelsson et al., 2007). The Phobius web server (Käll et al., 2007) was used to predict transmembrane domains for all proteins.

### Literature references

Literature references for characterized proteins were assigned to FOG’s in AYbRAH. Additional references were obtained from paperBLAST (Price and Arkin, 2017), UniProt (Consortium, 2014), *Saccharomyces* Genome Database (Cherry et al., 2011), PomBase (McDowall et al., 2014), *Candida* Genome Database (Inglis et al., 2011), and *Aspergillus* Genome Database (Cerqueira et al., 2013).

## 3 AYBRAH OVERVIEW

AYbRAH v0.1 and v0.2.3 database statistics are summarized in Table 2. In total, there are 214 498 protein sequences in the pan-genome for 33 yeasts and fungi; Pezizomycotina fungi were included in the database as an outgroup because they have genes that were present in Proto-Yeast’s ancestor, but subsequently lost. AYbRAH has 187 555 proteins (87% of the pan-proteome) that were assigned to 22 538 ortholog groups, and 18 202 homolog groups. Ortholog assignments are available in Excel, a tab-separated file, orthoXML (Schmitt et al., 2011), and a JSON format.

**Table 2.** AYbRAH ortholog database statistics before and after curation. The initial ortholog assignments were obtained with OrthoMCL and OrthoDB. Additional proteins were annotated using TBLASTN. Ortholog groups for enzymes and small metabolite transporters were manually curated by visual inspection of homolog phylogeny, and by identifying ortholog groups with an ETE 3 script (Huerta-Cepas et al., 2016a). Ortholog groups were modified by adding proteins to existing groups via pplacer (Matsen et al., 2010), or by collapsing homolog groups into a single ortholog group if there were no gene duplications in the homolog group (under-clustering).

|  | AYbRAH |  |
| --- | --- | --- |
|  | v0.1 | v0.2.3 |
| Proteins | 212 551 | 214 498 |
| Proteins in AYbRAH | 169 118 (79%) | 187 555 (87%) |
| Ortholog groups | 14 249 | 22 538 |
| Homolog groups | 0 | 18 202 |
| Manually curated ortholog groups | 0 | 625 |
| Electronically modified ortholog groups | 0 | 3760 |

Ortholog groups are considered to be equal or siblings in current ortholog databases and COG collections (Galperin et al., 2017). AYbRAH adopts multi-level hierarchical relationships between ortholog groups, which was recommended by Galperin et al. (2017). An example is illustrated in Figure 3 for the acetyl-CoA synthetase homolog group (HOG00229). Acs1p was present in the common ancestor of all yeasts and fungi, but a duplication of *ACS1* led to *ACS2* in budding yeasts. Therefore, Acs1p (FOG00404) is the parent ortholog group to Acs2p (FOG00405). These relationships can be mapped to a species tree to illustrate how function has been gained and lost.

**Figure 3.**
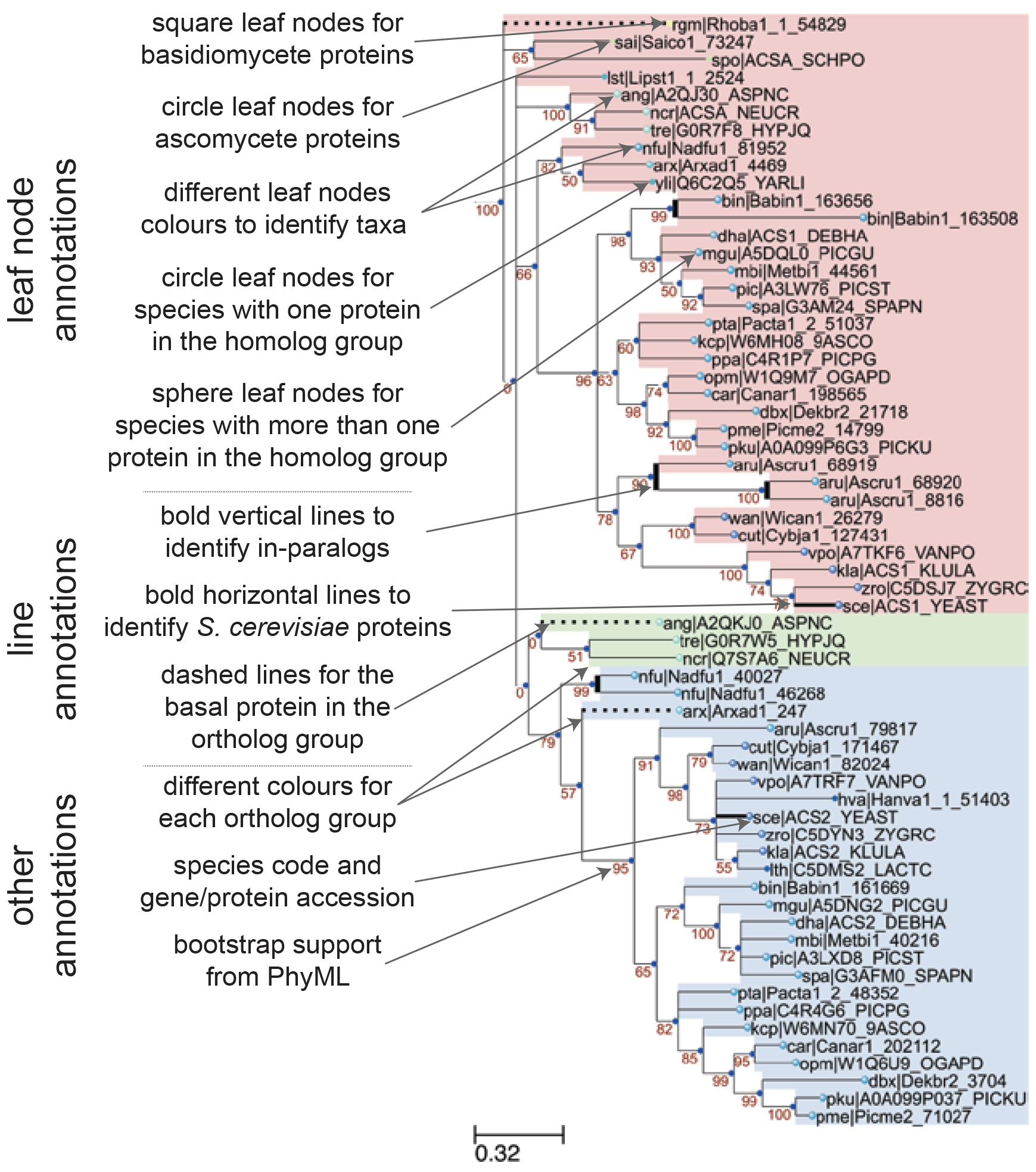
Annotation features of a sample phylogenetic tree in AYbRAH. Square and circle leaves indicate protein sequences in Basidiomycota or Ascomycota, respectively. Leaf nodes are coloured based on taxonomic groups. Circle leaves are used for proteins with no paralogs in the same species, whereas sphere leaves are used to designate proteins with paralogs in the same species. Vertical bold lines indicate species-lineage expansions, which are sometimes called in-paralogs or co-orthologs (Remm et al., 2001). Horizontal bold lines designate *S. cerevisiae* proteins, which is the most widely studied eukaryote. Dashed lines indicate the most anciently diverged protein sequence in the ortholog group. Ortholog groups can be identified by colour groups to help the visual inspection of ortholog assignments. The leaf names include a three-letter species code and a sequence accession. Internal nodes are labelled with the bootstrap values from phylogenetic reconstruction with PhyML.

### The AYbRAH web portal

AYbRAH has a web page for each homolog group with information on gene names, descriptions, gene origin (paralog, ohnolog, xenolog), literature references, localization predictions, and phylogenetic reconstruction. Homolog groups can be searched by FOG (FOG00404) or HOG (HOG00229) identification codes, gene names (*ACS1*), ordered locus (YAL054C), UniProt entry names (ACS1 YEAST), or protein accession codes from UniProt (Q01574), NCBI RefSeq (NP 009347.1) or EMBL (CAA47054.1).

Snapshots for mitochondrial localization and transmembrane domain predictions are shown in Figures 4A and 4B for internal alternative NADH dehydrogenase, encoded by *NDI1* (FOG00846). Reviewing localization predictions for orthologous proteins with multiple algorithms enables researchers to make prudent decisions about protein localization, rather than relying on one method for one protein sequence. For example, Cybja1 131289 encodes internal alternative NADH dehydrogenase, yet its mitochondrial localization probability is 0.0019 with MitoProt II; all other mitochondrial predictions for Ndi1p orthologs are greater than 0.80 with MitoProt II. A review of the upstream nucleotide sequence of Cybja1 131289 indicates additional start codons that were not included in the protein annotation. MitoProt II predicts a mitochondrial localization probability of 0.5191 for the full protein sequence, which is more consistent with its orthologs.

**Figure 4.**
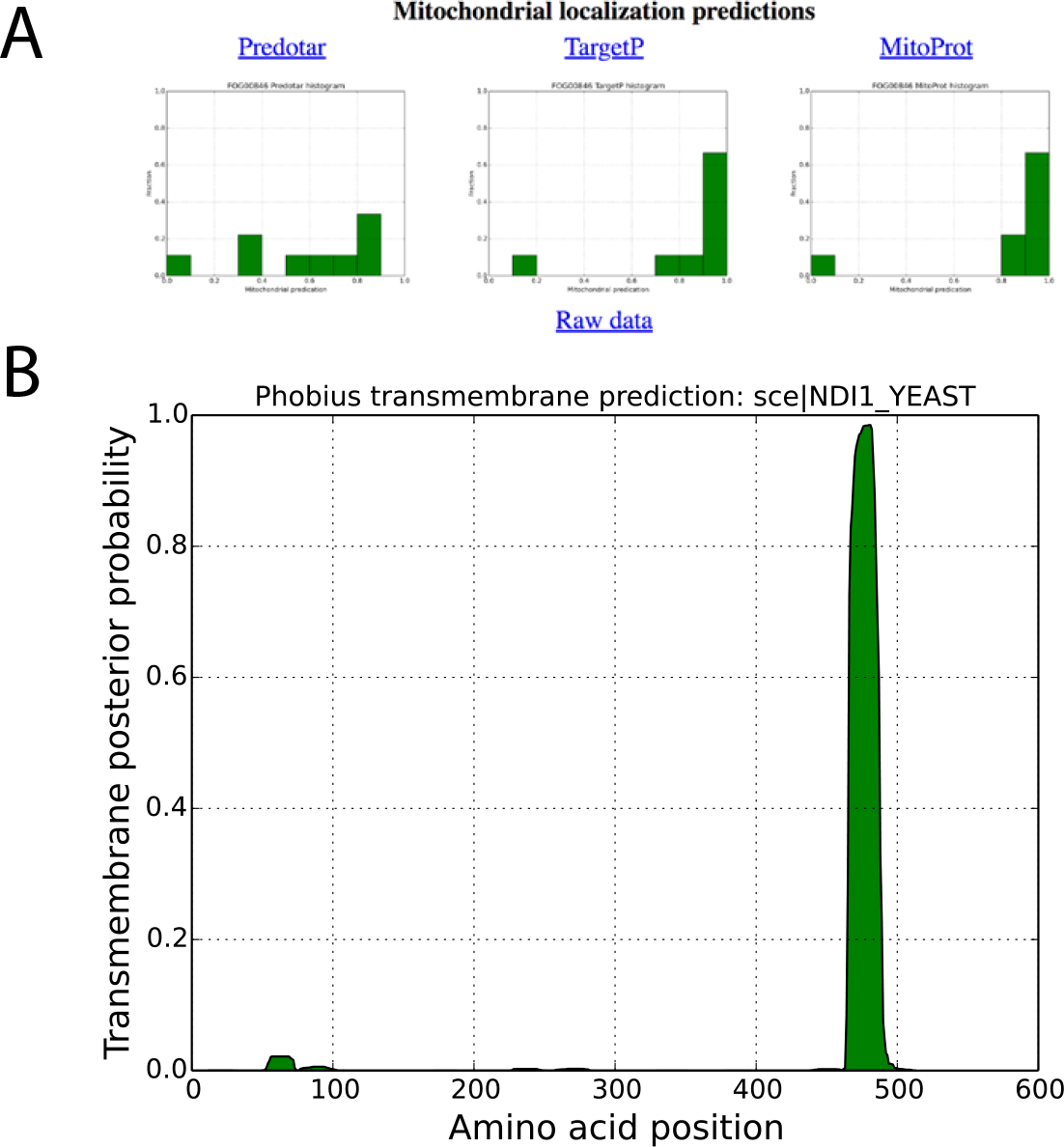
Localization predictions for internal NADH dehydrogenase (NDI1 YEAST) in AYbRAH. (A) Histogram plots are shown for mitochondrial localization predictions of orthologous Ndi1p predicted with Predotar, TargetP, and MitoProt. (B) Transmembrane domain predictions computed for orthologous proteins by the Phobius web server.

A sample phylogenetic tree rendered by ETE v3 (Huerta-Cepas et al., 2016a) and descriptions of its annotation features is shown in Figure 3 for the acetyl-CoA synthetase family (HOG00229). The initial ortholog assignment by OrthoMCL did not distinguish between the *Acs1p* (FOG00404) and *Acs2p* (FOG00405) paralogs. Discrepancies in ortholog assignments can be identified by comparing bootstrap support values for clades with the assigned ortholog groups; issues can reported on GitHub or issuing pull requests for large changes to ortholog groups.

## 4 AYBRAH CURATION

OrthoMCL and OrthoDB are less computationally intensive than phylogenetic-based methods but are not always accurate (Salichos and Rokas, 2011). Curation was required to resolve incorrect ortholog assignments due to over-clustering and under-clustering.

### Over-clustering by OrthoMCL

Over-clustering has been described in past studies (Jothi et al., 2006), which occurs when graph-based methods create ortholog groups that do not distinguish between orthologs and paralogs. Over-clustering by OrthoMCL was common in gene families with many duplications or high sequence similarities, such as the aldehyde dehydrogenase (HOG00216) and the major facilitator superfamily (HOG01031); adjusting parameters for BLASTP and OrthoMCL did not help differentiate between orthologs and paralogs in HOG00216 (results not shown). Figure 5 illustrates an example of over-clustering with a subset of the hexokinase family (HOG00193). In this phylogenetic reconstruction, one hexokinase gene is present in the ancestral yeast species, but a gene duplication in Pichiaceae leads to the *HXK3* paralog; the *HXK2* ortholog is subsequently not maintained in *O. parapolymorpha*’s genome. OrthoMCL assigned the *HXK3* paralog to the same ortholog group as *HXK2*. The bidirectional best hit method, commonly used for ortholog identification (Moreno-Hagelsieb and Latimer, 2008), would have also falsely identified *O. parapolymorpha*’s *HXK3* as orthologous to *S. cerevisiae*’s *HXK2*. This example highlights how the greediness of graph-based methods can misidentify orthologs, which has been shown for yeast ohnologs (Salichos and Rokas, 2011), and how incorrect ortholog assignments can be made with pairwise comparisons. Paralogs were identified from over-clustered ortholog groups by finding nodes with high bootstrap support in the the consensus phylogenetic trees for homologs and migrating the proteins to a new ortholog groups; in some cases orthologs were identified by reviewing the sequence alignment of homologs.

**Figure 5.**
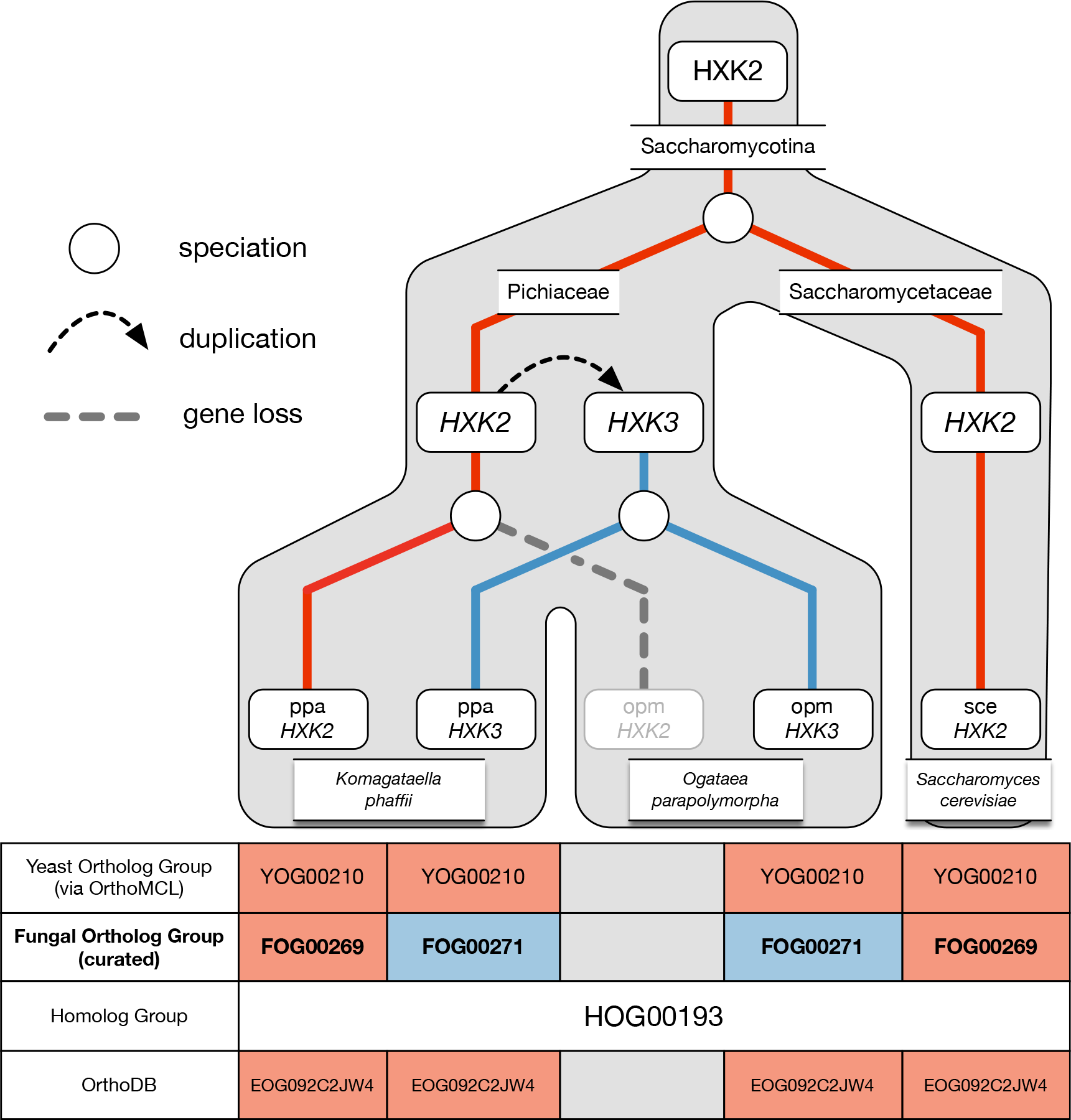
Example of over-clustering by OrthoMCL with the hexokinase family and its curation in AYbRAH. A gene duplication of *HXK2* in Pichiaceae led to the *HXK3* paralog. *HXK2* was subsequently lost in *Ogataea parapolymorpha* but maintained in *Komagataella phaffii*. OrthoMCL was unable to differentiate between the *Hxk2p* and *Hxk3p* orthologs. Both ortholog groups are also assigned to the same Fungi-level ortholog group in OrthoDB.

### Under-clustering by OrthoMCL

Under-clustering occurrs when orthologous proteins are assigned to multiple ortholog groups.

OrthoMCL was more prone to under-clustering by short protein sequences and protein sequences with low sequence similarity, such as electron transport chain subunits and Flo8p. Figure 6 demonstrates under-clustering with a subset of the Flo8p family that was not assigned to one ortholog group by OrthoMCL. Under-clustering was mostly resolved via a Python script that coalesced proteins into a new ortholog group when multiple ortholog groups were present in the homolog group yet no organism had any gene duplications.

**Figure 6.**
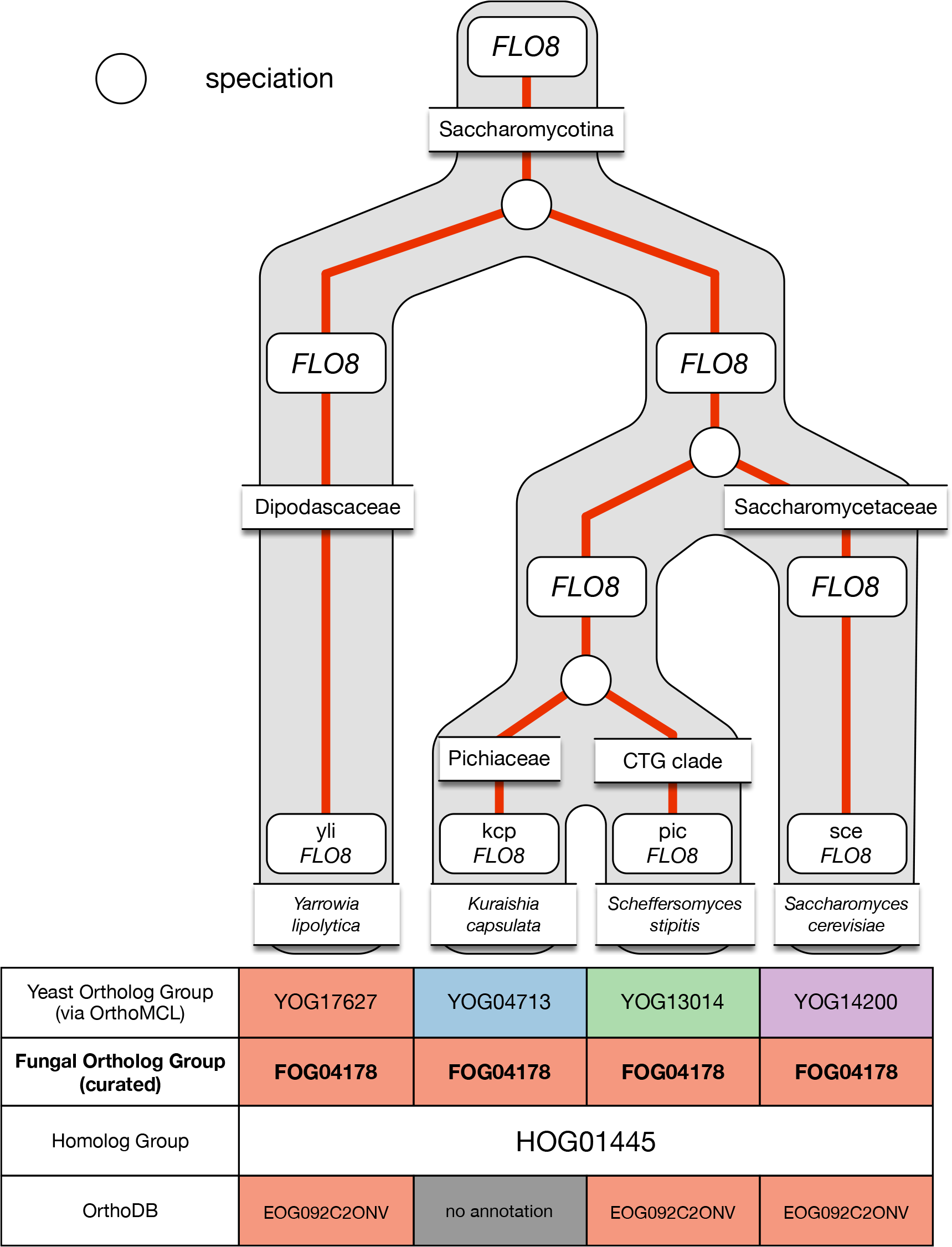
Example of under-clustering by OrthoMCL in the *FLO8* ortholog group and its curation in AYbRAH. OrthoMCL dispersed the Flo8p proteins into multiple ortholog groups due to the low sequence similarity between the proteins. The proteins were merged into one ortholog group.

## 5 COMPARISON OF AYBRAH TO OTHER ORTHOLOG IDENTIFICATION METHODS

### BLASTP scoring metrics

BLASTP is used as the basis for many ortholog predictions, including graph-based methods (Kuzniar et al., 2008) and reciprocal best hits (Moreno-Hagelsieb and Latimer, 2008). The distribution of percent identity, log(bitscore), and −log(expect-value) for proteins identified as orthologous to *Saccharomyces cerevisiae*’s proteins in AYbRAH are shown in Figure 7. Taxonomic groups include the Saccharomycotina outgroup, basal Saccharomycotina, Pichiaceae, the CTG clade, Phaffomycetaceae and Saccharomycodaceae, and Saccharomycetaceae (Table 1). The approximate divergence time with *Saccharomyces cerevisiae* is 400-600 million years with the Saccharomycotina outgroup, 200-400 million years with the basal Saccharomycotina yeasts, 200 million years with Pichiaceae and CTG clades, 100-200 million years with Phaffomycetaceae and Saccharomycodaceae, and 0-100 million years with Saccharomycetaceae. The distributions of percent identity, log(bitscore), and −log(expect-value) for proteins with 100-400 million years of divergence with *S. cerevisiae* are similar; however, the distributions skew differently for percent identity and −log(expect-value) for the Saccharomycotina outgroup (more than 400 million years of divergence) and Saccharomycetaceae (less than 100 million years of divergence). Distributions for percent identity, log(bitscore), and −log(expect-value) for each species in AYbRAH are shown in Figures S1, S2, and S3. These results highlight the need to use phylogenetic methods and hidden Markov models to identify orthologs over long evolutionary timescales (Mi et al., 2016), but also enable orthologs to be identified by synteny and sequence similarity over smaller evolutionary time ranges (Byrne and Wolfe, 2005; Scannell et al., 2006).

**Figure 7.**
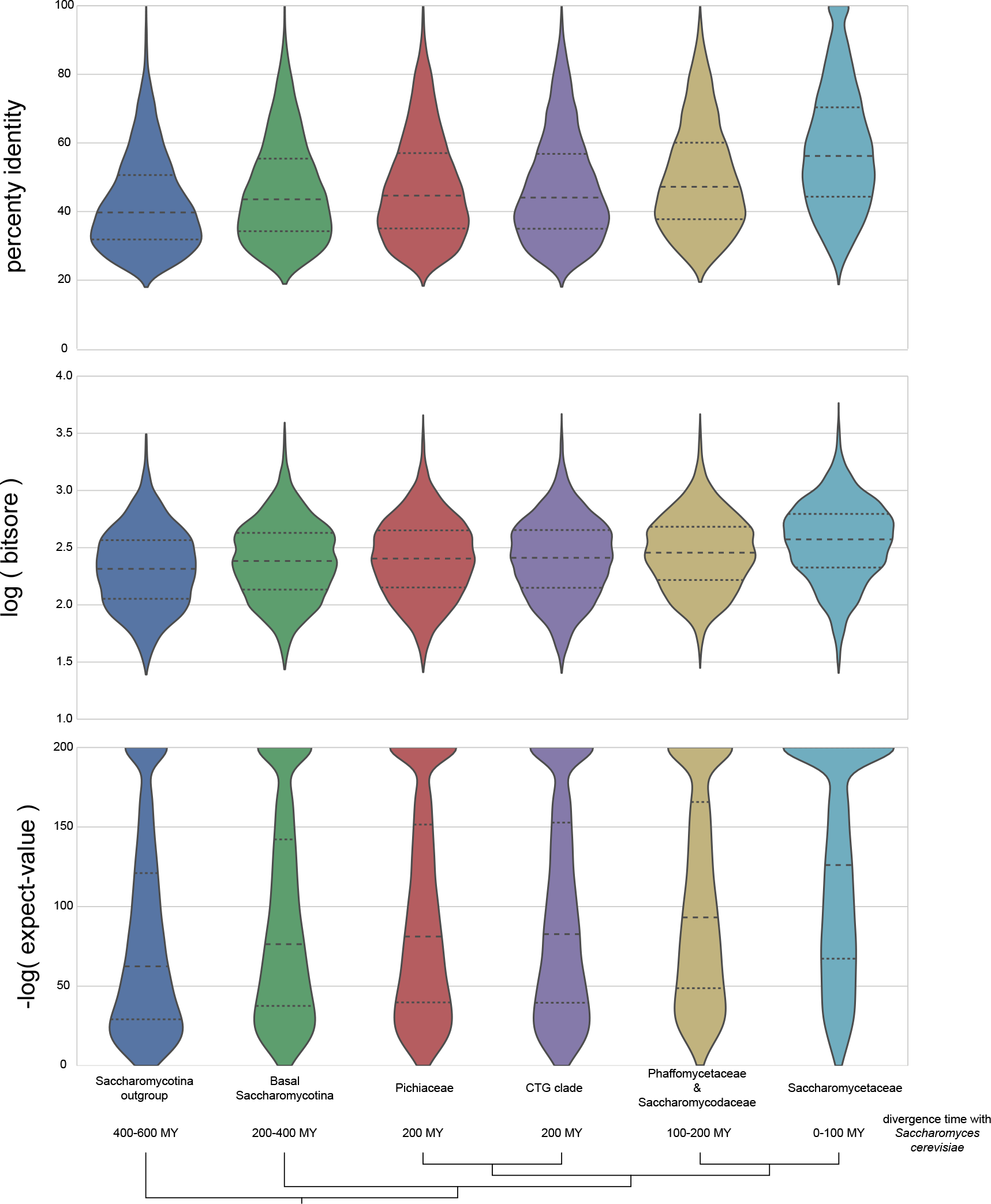
Comparison of BLASTP percent identities, logarithm of bit scores, and negative logarithm of expect-values for proteins orthologous to *Saccharomyces cerevisiae*. The bottom half of orthologous proteins in the Saccharomycotina outgroup and Saccharomycetaceae have a percent identities of less than 40% and 58%, respectively; the bottom half of the expect-value ranges is more than 1e-60 and 1e-125 for the same groups. The wide and skewed distribution in the Saccharomycotina outgroup highlights the difficulty in making pairwise ortholog predictions for proteins with more than 400 millions of divergence in Dikarya fungi with BLASTP results; however orthologs can be easily identified in the Saccharomycetaceae family because of their high sequence similarities and low expect-values.

### Comparison of AYbRAH and well-established phylogenomic databases

Ortholog assignments in AYbRAH were compared with OMA, PANTHER, HOGENOM, eggNOG, and KO (Table 3). OMA and PANTHER have the highest number of congruous ortholog groups with AYbRAH. Interestingly, PANTHER tends to over-cluster protein sequences into ortholog groups, while OMA tends to under-cluster. HOGENOM, eggNOG, and KO have a high fraction of proteins not assigned to any ortholog groups.

**Table 3.**
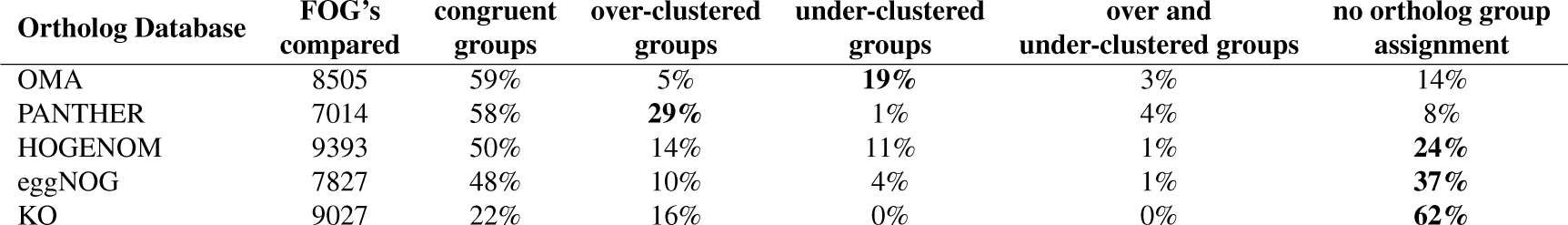
Comparison of ortholog assignments between AYbRAH and well-established phylogenomic databases. OMA and PANTHER are the most congruous with AYbRAH; OMA and PANTHER are predicted to have more under-clustered and over-clustered groups relative to AYbRAH, respectively. HOGENOM, eggNOG, and KO have a large number of proteins with no ortholog assignment.

| Ortholog Database | FOG's compared | congruent groups | over-clustered groups | under-clustered groups | over and under-clustered groups | no ortholog group assignment |
| --- | --- | --- | --- | --- | --- | --- |
| OMA | 8505 | 59% | 5% | 19% | 3% | 14% |
| PANTHER | 7014 | 58% | 29% | 1% | 4% | 8% |
| HOGENOM | 9393 | 50% | 14% | 11% | 1% | 24% |
| eggNOG | 7827 | 48% | 10% | 4% | 1% | 37% |
| KO | 9027 | 22% | 16% | 0% | 0% | 62% |

10 ortholog groups were randomly selected from the over-clustered groups in PANTHER and under-clustered groups in OMA to determine the source of the incongruency. It was found that three of the 10 over-clustered ortholog groups in PANTHER were correctly annotated in AYbRAH, one ortholog group was correctly identified in PANTHER but under-clustered in AYbRAH, one ortholog group was not correctly identified in either database, and five ortholog groups required further curation since the phylogenies are ambiguous. All 10 ortholog groups from OMA were under-clustered, suggesting a systematic bias to not cluster proteins with lower sequence similarity; i.e., proteins identified as orthologous in AYbRAH were separated into two or more ortholog groups in OMA. Therefore, the PANTHER database is most closely aligned with AYbRAH. All other databases appear to be more prone to over-clustering, or not have any annotation.

## 6 APPLICATIONS OF A CURATED ORTHOLOG DATABASE

Ortholog databases offer additional benefits beyond identifying orthologous proteins. These databases can be used to identify gene targets for functional characterization to functional genome annotation to streamlining genome-scale network reconstruction; Galperin et al. (2017) recently outlined some of the benefits and challenges to ortholog databases for microbial genomics. First, a curated ortholog database can serve as a repository for orthologs that have been screened and orthologs that require screening (Galperin and Koonin, 2004). Rather than characterizing all the orthologs in a handful of model organisms, research communities can broaden their efforts to understand the orthologs that do not exist in model organisms and the set of orthologs that do not have a conserved function with orthologs in model organisms. Second, a curated ortholog database can be used to improve and simplify genome annotation (Galperin and Koonin, 2004). Genes from newly sequenced organisms can be mapped to curated ortholog groups, similar to eggNOG-mapper (Huerta-Cepas et al., 2017), rather than using protein sequences from ortholog databases as queries in TBLASTN searches (Proux-Wéra et al., 2012). New ortholog groups can be created for *de novo* genes or genes from recent duplications. Pulling annotations from a curated ortholog database has the advantage of unifying the names and descriptions of genes between organisms, as has been proposed for ribosomal subunits (Ban et al., 2014), and can reduce the number of genes that are misannotated or annotated as conserved hypothetical proteins. And finally, a curated ortholog database can be used to improve the quality and quantity of genome-scale network reconstructions (GENRE’s). GENRE’s inherently require a great deal of curation for ortholog assignment and functional annotation, which is often not transparent. Refocusing this effort to curate ortholog groups and their function in open-source knowledgebase for pan-genomes can allow for improvements to be pushed to all GENRE’s, and for GENRE’s to be compiled for any taxonomic level, from Kingdom to strain.

## 7 FUTURE PLANS FOR AYBRAH

### Integration with PANTHER

OrthoDB was chosen to cluster ortholog groups in AYbRAH into homolog groups because it spans more taxa than other phylogenomic databases, and has ortholog assignments for different taxonomic ranks; however, it is less specific than PANTHER, despite only having a few fungal proteome annotations. Future updates to AYbRAH will migrate the AYbRAH homolog group backbone from OrthoDB to PANTHER, and add the remaining fungi in PANTHER to increase its phylogenomic span. These include other fungal model organisms, fungi and yeasts having pathogenicity to humans or plants, or fungi and yeasts occupying important taxonomic ranks: *Batrachochytrium dendrobatidis*, *Cryptococcus neoformans*, *Puccinia graminis*, *Ustilago maydis*, *Emericella nidulans*, *Neosartorya fumigata*, *Phaeosphaeria nodorum*, *Sclerotinia sclerotiorum*, *Candida albicans*, and *Eremothecium gossypii*.

### Reconciling AYbRAH with YGOB and CGOB

YGOB (Byrne and Wolfe, 2005) and CGOB (Maguire et al., 2013) are the gold-standard for ortholog databases in yeast genomics, and were created using sequence similarity and synteny. YGOB and CGOB span roughly 112 and 239 million years of evolution, respectively, while AYbRAH spans more than 600 million years of evolution (Hedges et al., 2015). Although AYbRAH has a broader pan-genomic coverage, YGOB and CGOB are expected to have better paralog and ohnolog assignments than AYbRAH because of its use of synteny. Future versions of AYbRAH will be reconciled with YGOB and CGOB.

### Coordinate-based protein annotations

It has been noted that genome protein annotations sometimes contain inaccuracies (Maguire et al., 2013). For example, *Spathaspora passildarium*’s genome encodes two *PHO3* homologs (Correia et al., 2018), but only one protein is currently annotated. AYbRAH will adopt the genomic coordinate-based system used in YGOB and CGOB (Maguire et al., 2013) to improve protein annotations.

## 8 RECOMMENDATIONS FOR OTHER COMMUNITY-DRIVEN ORTHOLOG DATABASES

### Combing top-down and bottom-up approaches to identifying orthologs

In our experience PAN-THER and YGOB are the highest quality ortholog databases for yeasts. PANTHER spans some four billion years of evolution, has the most accurate ortholog assignments for large-scale phylogenomic databases when compared to AYbRAH, but has limited proteomes and cannot distinguish between recently emerged paralogs and ohnologs. On the other hand, YGOB spans 100 million years of evolution, also covers limited proteomes, but has high accuracy for recently evolved paralogs and ohnologs in Saccharomycetaceae. An alternative approach to curated ortholog databases for other taxa can leverage both methods. First, a research community can select a group of organisms of interest, and choose several proteomes in PANTHER that span the phylogenomic range. Next, TreeGrafter (Tang et al., 2018), or other phylogenetic placement algorithms, can be used to map the proteomes of interest to PANTHER families/subfamilies. Phylogenetic reconstruction of the proteins mapped to the PANTHER families or subfamilies can be used to distinguish any paralogs using tree-crawling scripts, such as ETE (Huerta-Cepas et al., 2016a), or by visual inspection. Unresolved paralog or ohnolog relationships can also be resolved using genomic synteny of closely related organisms, similar to YGOB (Byrne and Wolfe, 2005). This approach leverages the accuracy of PANTHER from the top-down for fungal genomes, and from the bottom-up using the genomic synteny from YGOB.

### Multi-level phylogenetic placement

Czech et al. (2018) recently outlined an approach to automating multilevel phylogenetic placement, which is often used to map environmental sequences onto an existing species tree reconstructions. This method can be adapted to ortholog identification with TreeGrafter (Tang et al., 2018) or pplacer (Matsen et al., 2010). Two sets of phylogenetic trees can be created in this multi-level approach. Smaller reference trees trimmed from phylogenetic trees of homologs that maintain sequence diversity of orthologs, and clade phylogenetic trees with higher sequence resolution. Proteins can then be queried against the reference tree and a subsequent clade tree with higher sequence resolution.

### Alternative phylogenetic programs

Some phylogenetic trees had low bootstrap support for ortholog groups, which prohibited identification of orthologs and paralogs. For example, NADP-dependent serine dehydrogenase (HOG00233) has several closely related ortholog groups. Additional programs for maximum likelihood-based phylogenetic analysis can be used to reconstruct the phylogeny of homologs, such as IQ-TREE (Nguyen et al., 2014), which has been shown to outperform RAxML and PHYML for species tree reconstruction (Zhou et al., 2017). Small phylogenetic trees can be reconstructed with subfamilies of the homolog groups.

## 9 CONCLUSION

In conclusion, we developed AYbRAH as an open-source ortholog database for yeasts and fungi. Manual curation was required for gene families with high sequence similarity, often arising from recent gene duplications, and with gene families with low sequence similarity. Curated ortholog databases can be implemented for other taxa to improve their genome annotations, or use them as benchmarks for orthology assignment methods.

## 10 ACKNOWLEDGEMENTS

The authors gratefully acknowledge Prof. Belinda Chang and Ryan Schott for their advice with the phylogenetic analysis, and Dean Robson for his help implementing the search function in the AYbRAH web portal.

## 11 AUTHOR CONTRIBUTIONS

KC and SM performed the phylogenetic analysis. KC and RM discussed the results. KC wrote the manuscript. KC and RM reviewed the manuscript.

## 13 ABBREVIATIONS

AYbRAH: Analyzing Yeasts by Reconstructing Ancestry of Homologs
CGOB: *Candida* Gene Order Browser
COG: Clusters of Orthologous Groups
FOG: Fungal Ortholog Group
HOG: Homolog Group
YGOB: Yeast Gene Order Browser

## 14 AVAILABILITY OF DATA AND MATERIALS

AYbRAH database files and additional files, such as phylogenetic trees and sequence alignments, can be found at https://github.com/kcorreia/aybrah.

